# Temperature-dependent Thermodynamic and Photophysical Properties of SYTO-13 Dye Bound to DNA

**DOI:** 10.1101/2022.08.31.505902

**Authors:** Robert F. DeJaco, Jacob M. Majikes, J. Alexander Liddle, Anthony J. Kearsley

## Abstract

The benefits of dyes in nucleic acid assays above room temperature are limited by a nonlinear, highdimensional relationship between fluorescence and the biophysical and chemical processes occurring in solution. To overcome these limitations, we identify an experimental regime that eliminates bias and unnecessary complexities in this relationship, and develop an experimental–computational workflow to generate the property data required to describe the dependence of fluorescence on temperature and concentration. Specifically, we exploit the temperature-cycling capabilities of real-time PCR machine, as well as the utility of numerical optimization, to determine the binding strength and molar fluorescence of the SYTO-13 dye bound to double-stranded (DS) or single-stranded (SS) DNA at more than 60 temperatures. We find that the data analysis approach is robust; it can account for significant well-to-well and plate-to-plate variation. The weak binding strength of SYTO-13 relative to SYBR Green I is consistent with previous reports of its negligible influence on PCR and melting temperature. Discriminating between molar fluorescence and binding strength clarifies the mechanism for the larger fluorescence of a DS/dye solution than a SS/dye solution; in fact, the explanation is different at high temperature than at low temperature. The temperature-dependence of the binding strength allows for ascertainment of the enthalpic and entropic contributions to the free energy, as well as the sign of the differential heat capacity of binding. The temperature-dependence of the molar fluorescence allows for calculation of the brightness (quantum yield times molar extinction coefficient) of SYTO-13 bound to DS relative to SS. The more accurate and complete description of the relationship between solution behavior and fluorescence enabled by this work can lead to more accurate selection of dyes and quantification of nucleic acids.

**SIGNIFICANCE:** Fluorescent dyes are often used to quantify nucleic acids. The accuracy and precision of quantification, however, is limited by a complex and high-dimensional relationship between fluorescence and solution behavior. This is especially true for assays above room temperature, where empirical approximations are often required in the absence of available property data. In this work, we present an experimental and computational workflow that can more accurately describe this relationship and more efficiently generate the temperature-dependent thermodynamic and photophysical properties required. This approach can improve quantification and selection of high-performing dyes for particular assays.

## INTRODUCTION

A wide variety of nucleic acid assays employ fluorescent probes to measure a signal that can be associated with a specific biological phenomenon. Organic dyes that interact with nucleic acids(1) are significantly less expensive than covalently attached probes. They can also be readily used in different assays, as they are not designed for a specific sequence.

However, their less specific nature limits their quantitative utility. In contrast to fluorescence associated with covalently attached probes that bind strongly to their compliment, the fluorescence of dyes is intimately related to the partitioning of dye between DNA and solution. As a result, an additional property, the partition coefficient, is required to related solution behavior to measured fluorescence.

The influence of binding equilibria on fluorescence is routinely addressed in spectrofluorimetry(2–8). In fact, spectrofluorimetry is typically used to extract equilibria from fluorescence measurements. However, many studies attribute nonlinearities in fluorescence to packing effects; such behavior may also arise from dimerization in solution(9) and self-quenching(10). At the same time, the sources and magnitudes of error in the parameter fitting procedure are typically not discussed. This is important to aid in comparing results from different laboratories that are sometimes conflicting (see Table 2).

Unlike spectrofluorimetric techniques, real-time PCR and high-resolution melting assays do not usually account for the influence of dye binding equilibria on fluorescence. This is likely due to a lack of available property data above room temperature. The conventional expressions used for each assay turn out to result from different approximations (see Materials and Methods). A greater availability of temperature-dependent binding equilibria could enable the validity of each assumption to be assessed or relaxed entirely.

In this work, we present a new technique for extracting temperature-dependent partition coefficients for a dye binding to double-stranded DNA (DS) or single-stranded DNA (SS) from a thermocycler instrument measuring fluorescence. Careful attention to the mathematical model reveals that, by focusing on an experimental regime where the dye concentration is low enough for the fluorescence to be linear in dye concentration, bias due to quenching or deviations from Beer’s Law can be eliminated. Analysis of the model also reveals that an additional property, the molar fluorescence (the fluorescence analogue of the extinction coefficient), can also be extracted at each temperature.

Subsequently, experiments are undertaken to apply the method to the SYTO-13 dye interacting with DS or SS. However, the experimental data exhibits noise both within a 96well plate and between replicate plates. A conventional approach of minimizing the error in fluorescence signal yields unphysical interpretations of plate-to-plate variation. It suggests that the instrument underestimates the measured signal on one day, but overestimates on another. Accounting for error in total dye concentration (in addition to error in signal) yields errors that are consistent with the pipetting procedure and a machine that is routinely calibrated. Such a quantitative understanding of error aids in reproducibility.

In addition to generating a large amount of thermodynamic and photophysical property data in a robust and efficient manner, the approach provides useful insight into the behavior of SYTO-13 and DNA. The relatively weaking binding of SYTO-13 explains its minimal impact on PCR and melting temperature. Deconvoluting the brightness and binding strength provides a mechanistic explanation for the larger total fluorescence of DS than SS. At low temperature, the fluorencence of DS is larger due to the larger binding strength and molar fluorescence. At higher temperature, the fluorencence associated with one dye bound to DS is very similar to that of one dye bound to SS. In this case, the larger total fluorescence of DS is due to more dye adsorbing per nucleotide.

## MATERIALS AND METHODS

### Mathematical Model

The dependence of the fluorescence *F* = *F*(*C, D, T*) on total dye concentration, *C*, total DNA concentration, *D*, and temperature, *T*, may be described by

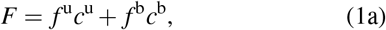

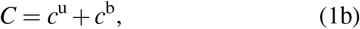

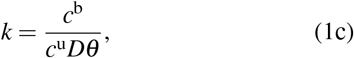

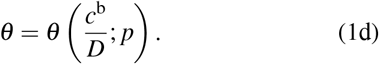

Here, *c*^u^ is the concentration of unbound dye, and *c*^b^ is the concentration of bound dye (or the concentration of the dye– DNA complex). When *F* corresponds to an absorbance measurement, *f* ^*i*^(*T*) = *𝓁ε*^*i*^(*T*) for each species *i*, where *𝓁* is the optical path length and *ε*^*i*^ is the molar extinction coefficient. In this case, (1a) corresponds to Beer’s Law for two attenuating species. When *F* corresponds to a fluorescence measurement, which is undertaken in this work, the quantity *f* ^*i*^, referred to as the fluorescence per mole of species *i*, represents(2, 11)

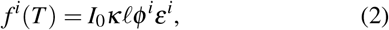

where *I*_0_(*T*) is the excitation intensity, *ϕ*^*i*^(*T*) is the quantum yield associated with species *i*, and *κ* is an instrument-specific constant depending on the collection efficiency of the emitted light. The quantity *ϕ*^*i*^*ε*^*i*^ is often called the brightness(12) of species *i*.

In Equation (1c), *k* = *k*(*T*) represents the equilibrium constant for the reaction between a dye molecule, a vacant adsorption site on DNA, and an adsorption site on DNA occupied by dye, or

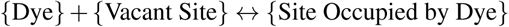

In this partitioning equation, *θ* represents the average number of free adsorption sites per strand of DNA. In (1d), a constitutive expression depending on a set of parameters *p* is chosen to describe the dependence of *θ* on *c*^b^*/D*, the average number of dye bound to DNA. The expression is very frequently taken from that of McGhee and von Hippel(13), although there are other appropriate options at low coverage(14).

Rearrangement and substitution of (1b), (1c), and (1d) into (1a) yields

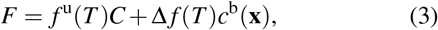

where Δ*f* = *f*^b^ *− f*^u^, and **x** is defined as the tuple

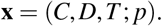

To understand the dependence of *c*^b^ on **x**, substitute (1d) and *c*^u^ = *C − c*^b^ from (1b) into (1c), and rearrange to obtain an equation *g*(*C, D, T, p, c*^b^) = 0. Since *θ* is a decreasing function of *c*^b^, the implicit function theorem demonstrates that *c*^b^ = *c*^b^(**x**).

Equation (3) is the conventional model used in spectrofluorimetry or spectrophotometry. With suitable experiments, the relation allows for fitting of the parameters *f*^b^, *f*^u^, *k*, and *p* to experimental data at each temperature *T*. However, this model assumes that *f*^b^ and *f*^u^ are independent of concentration. At high concentrations, deviations from Beer’s Law or quenching may be present. This means that *f*^*i*^ = *f*^*i*^(*c*^*i*^) for each species *i*. Since *c*^u^ = *C − c*^b^, (3) is really an approximation of

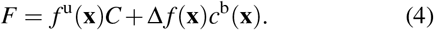

Equation (4) demonstrates why the conventional approach is biased at high dye concentrations. Equation (3) assumes that, for example, the fluorescence of 5 complexes of 1 dye bound to each DNA strand is identical to the fluorescence of 5 dye bound to 1 DNA strand (see Figure 1). The more generic Equation (4) demonstrates that, when *F* is a nonlinear function of concentration, the underlying biophysical processes may involve any combination of nonlinear packing effects (adsorption), deviations from Beer’s Law, or quenching. Quenching and deviations from Beer’s Law can occur both for dye in solution and dye bound to DNA.

**Figure 1:**
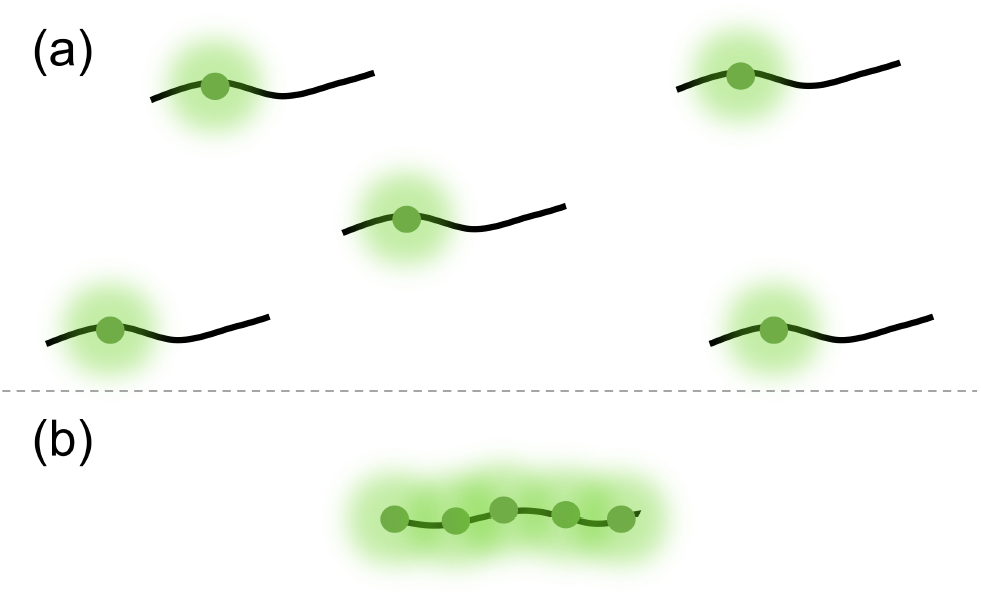
Visual depiction of fluorescence associated with (a) 5 complexes of 1 dye bound to a DNA strand and (b) 5 dye bound to 1 DNA strand. The conventional approach assumes that the fluorescence of (a) and (b) are equivalent, as *c*^b^ is the same.

Such complications can be alleviated at low dye concentrations where the fluorescence is a linear function of dye concentration. In this regime, the number of free adsorption sites is constant at *θ*_max_, or

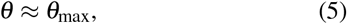

and, (1c) may be rearranged to

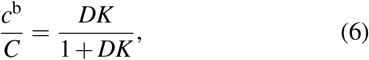

where *K* = *kθ*_max_. Equation (6) is an explicit expression for the fraction of dye bound to DNA. In this dilute regime, where Beer’s Law is valid and quenching is negligible,

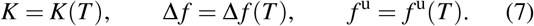

Substitution of (6) and (7) into (4) yields

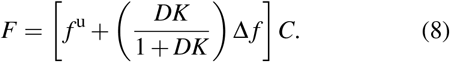

Equation (8) is a key result. It demonstrates that, in the dilute regime, *F* is a linear function of total dye concentration. Rather than making assumptions, assessing the degree of linearity of *F* with respect to *C* allows one to determine whether the model is valid. The validity of the approximation (5) can be further checked by assessing whether the average number of dye per DNA, or

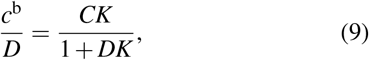

is small. Using experimental conditions that simplify the relationship between fluorescence and solution behavior facilitates the goal of using the reporter to learn more information about underlying characteristics of DNA.

The model used in this work, Equation (8), is a slight modification of the fluorescence analogue of the initial models used by Benesi and Hildebrand(15) and Peacocke and Skerrett(16) over 65 years ago. Yet, the impact of dye binding strength is typically not accounted for in real-time PCR and high-resolution melting. As such, it is useful to investigate what assumptions are being made in conventional techniques. In real-time PCR, the standard curve calculates amplification efficiencies by assuming that the fluorescence is linearly proportional to the total amount of DNA. From (8), we see that the linear dependence of *F* on *D* only occurs when *DK →* 0. This can only be true after significant DNA has amplified when *K →* 0, or when adsorption is very weak.

In a melting experiment, dye partitions between SS, DS, and solution. Equation (8) instead becomes

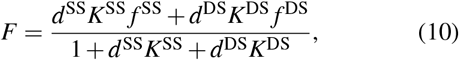

where the superscript b has been replaced with SS or DS, *d*^*i*^ corresponds to the concentration of DNA of type *i* on a singlestranded basis, and *K*^*i*^ denotes the partition coefficient for each type *i*. (Without loss of generality, we have assumed that *f*^u^ = 0.) Defining the fraction melted as *d*^SS^*/D*, accounting for the mass conservation of DNA via *D* = *d*^SS^ + *d*^DS^, and rearranging yields an expression for the fraction melted in terms of *F, D,C, K*^SS^(*T*), *f*^SS^(*T*), *K*^DS^(*T*), and *f*^DS^(*T*). This expression is a linear function of the fraction melted. Provided *K*^SS^ (*f*^SS^ *– F*) ≠ *K*^DS^ (*f* ^DS^ *– F*), it and can be solved to yield

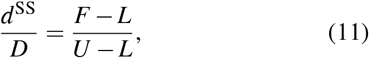

where *L* and *U* are defined as

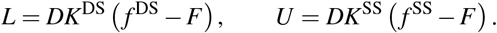

High-resolution melting analysis typically assumes that *L* and *U* are linear or exponential functions of temperature(17). A greater availability of temperature-dependent property data can assess the validity of these assumptions or relax them entirely.

### Experimentation

To determine a regime for which measured fluorescence is a linear function of dye concentration at constant DNA concentration and temperature, 96 well plates were prepared with dye concentrations increasing from left-to-right and top-to-bottom, corresponding to values ranging from 0.04 *μ*mol/L to 3.74 *μ*mol/L (see Table S1 in the Supporting Material, SM). In such a regime, (8) can be leveraged to calculate the partition coefficient *K* and molar fluorescence *f*^*p*^ of dye in each phase *p* at each temperature.

The SYTO-13 dye was selected for experimentation and obtained from Thermofisher Scientific, as previous work suggests that it has a negligible influence on DNA thermodynamics(18, 19). The DNA concentration, *D*, was kept the same in all wells of the plate at a value of 0 *μ*mol/L, 1.0 *μ*mol/L, or 2.0 *μ*mol/L. To keep track of different DNA concentrations, we will use subscript *k* = ⌊*D*⌋ (⌊ · ⌋ denoting the floor function) and refer to the value as *D*_*k*_. The nominal total volume of solution was 40 *μ*L for all wells and all plates. After sealing each plate with adhesive film, the liquid was vortexed and spun down to the bottom of the wells, and the plate was placed in a StepOnePlus real-time PCR machine^1^. A sequential temperature profile was programmed into the machine to collect data for the same plate at multiple temperatures. Specifically, the plate was heated to 363 K (90^*°*^C) for a 20 s denaturation step, rapidly cooled to 353 K (80^*°*^C), where data acquisition occurred after 4 min. The plate was then subjected to iterative cycles of 0.5 K cooling, 4 min equilibration, and data acquisition. That is, for each cycle *j* = 1 to 139, the temperature was

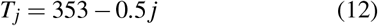

in units of Kelvin. (This corresponds to 80 *−* 0.5 *j* in units of Celsius.)

For each plate at each DNA concentration, two or three replicate plates, referred to as A, B, or C, were studied. As a control, we perfomed the set of experiments described above with two different types of DNA: (i) single-stranded DNA (SS) with sequence 5^*′*^-GACGAGGACGCAGCAGGCAGAC3^*′*^, and (ii) double-stranded DNA (DS) with both complementary strands. The concentrations of DS are defined so that *D* corresponds to the same number of nucleobases in solution as SS. That is, the concentration of each complementary strand of DS (or, equivalently, the concentration of the complex) is *D/*2. The fluorescence data was obtained from the GREEN channel.

To reflect the use of (8) with multiple experimental data points, the dye concentration *C* is hereafter viewed as

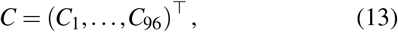

ordered so that *C*_*i*_ *>C*_*i−*1_ for each *i* ≥ 2. Similarly, the fluorescence data collected from thermocycling of each type of DNA *t* = SS or DS (or in the absence of DNA), each replicate plate *𝓁* =A, B, or C, and DNA concentration index *k* is denoted as

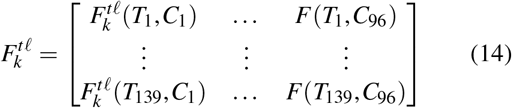

The tuple (*t, 𝓁, k*) thus distinguishes each plate from one another. It is used for naming in Table 2 and in the subplots of several figures.

## RESULTS AND DISCUSSION

### Raw Data and Scaling

The raw data are presented in Figure 2. Each row corresponds to a collection of replicate plates.

**Figure 2:**
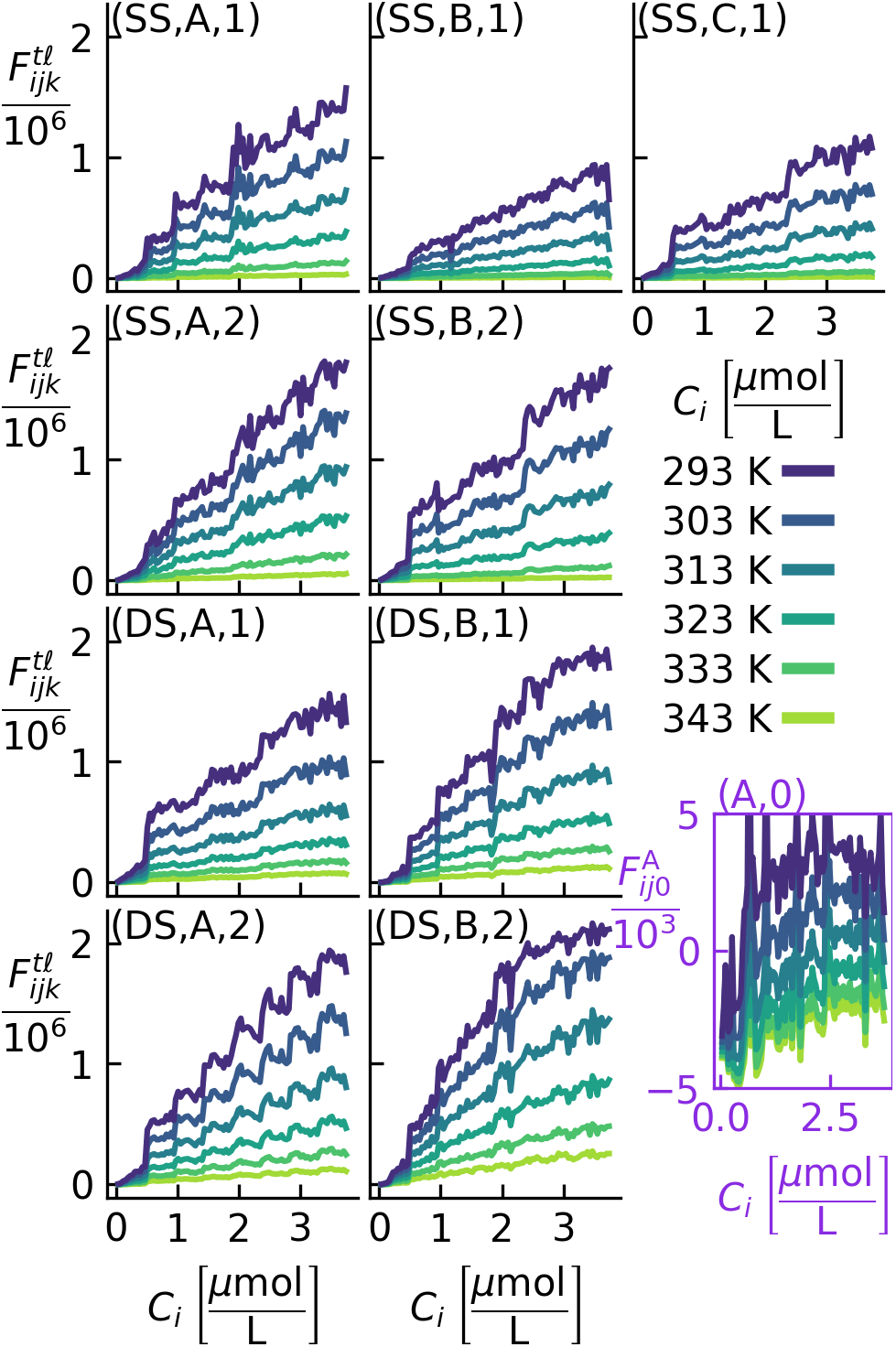
Visualization of fluorescence (vertical axes) as a function of dye concentration (horizontal axes) and temperature (color) for a variety of 96-well plates (subplots) in the presence of single-stranded DNA at *D* = 1 *μ*mol/L (first row), single-stranded DNA at *D* = 2 *μ*mol/L (second row), doublestranded DNA at *D* = 1 *μ*mol/L (third row), double-stranded DNA at *D* = 2 *μ*mol/L (fourth row), and without DNA (bottom right, purple axes). The dye concentrations in each plate are all the same, as depicted in Table S1 of the SM. The notation 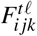 is an abbreviation for 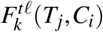, with the (*t, 𝓁, k*) tuple as named in Table 1 and used to label each subplot in the top left corner. For ease in visualization, only a few temperatures are shown.

**Figure 3:**
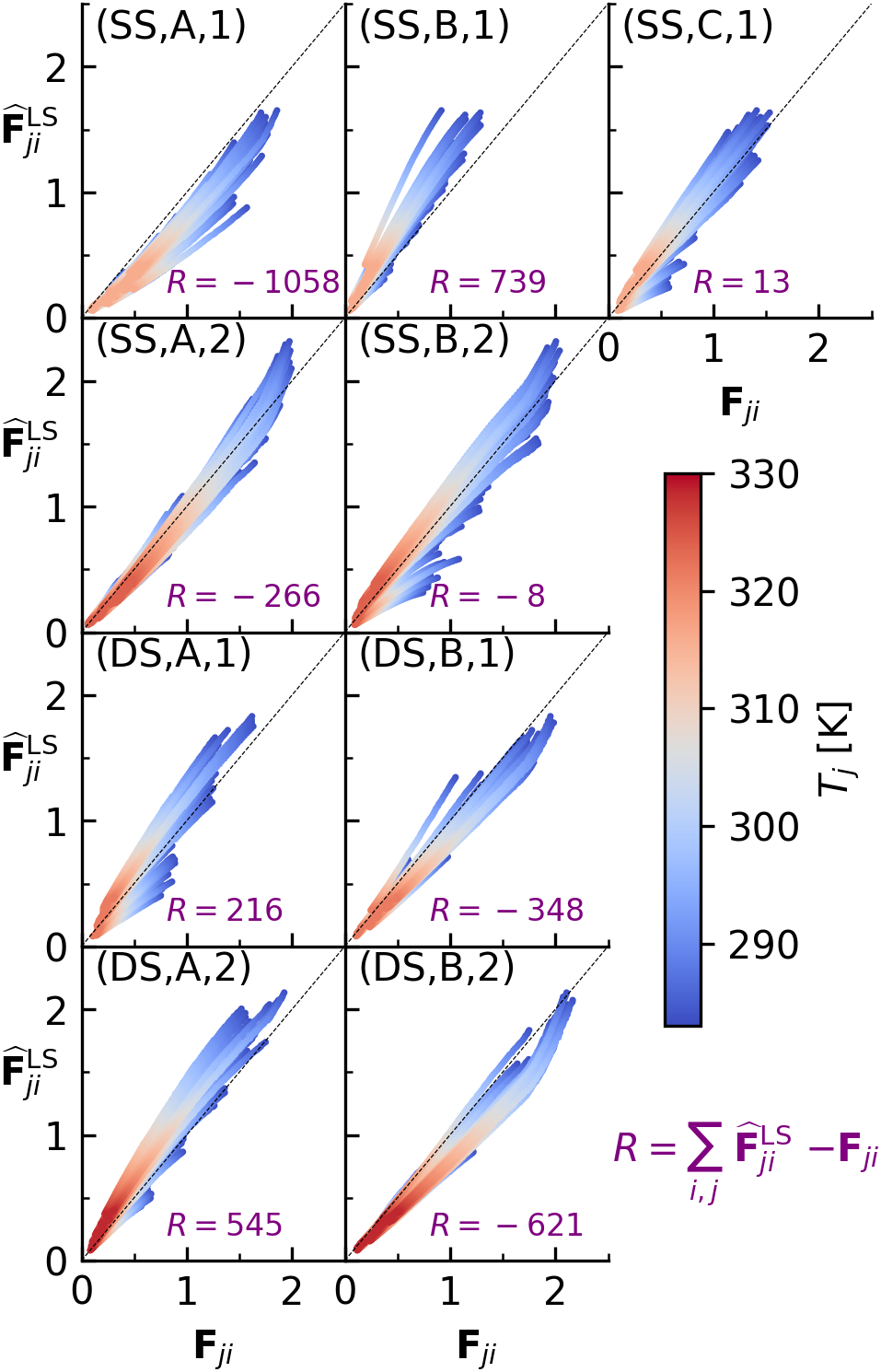
Fluorescence signal predicted from model, 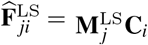, as a function of signal measured experimentally, or **F** _*ji*_, for each temperature index *j* and well *i*. Each subplot represents a distinct plate, with the label in the top left corresponding to Table 1. The colors correspond to the temperatures depicted in the colorbar. The colors are plotted at each coordinate with hot temperatures plotted on top of cold temperatures. The black dashed line indicates a location where 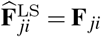, corresponding to perfect agreement. Data points below this line indicate an underestimation of the measured signal, while those above indicate an overestimation.

The bottom-right, purple-colored subplot in Figure 2 corresponds to a plate of wells that do not contain DNA. The vertical axis is scaled differently to reflect a significantly smaller signal. In this case, without DNA present, the fluorescence is very close to zero and sometimes negative. These unphysical values support the assumption that

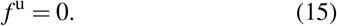

An analogous version of this assumption was also found to be appropriate in other work investigating the fluorescence for similar dyes interacting with DNA(3, 4, 7).

Wells possessing DNA appear to possess several common trends. First, there is significant well-to-well and plate-to-plate variation. For each replicate measurement at fixed temperature, the fluorescence often appears to oscillate around a line. These oscillations sometimes appear to possess a kind-of ‘sawtooth’ profile, with repetitive patters of dramatic increase followed by a series of small noisy changes. By comparing lines of the same color obtained by multiple replicate measurements, it can be observed that the underlying trends seem to be significantly different from one plate to another.

To account for well-to-well and plate-to-plate variation, we augment the collection of replicate plate data via

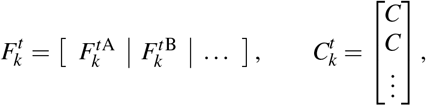

for each DNA type *t* and concentration *k*. The ellipses represent the additional entry that is only present for *t* = SS and *k* = 1, which has three replicates instead of two.

Second, all plates appear to have similar trends for dye concentrations less than 0.5 *μ*mol/L. Below this concentration, the fluorescence is near zero. As the dye concentration increases above 0.5 *μ*mol/L, the fluorescence exhibits a steep increase. This behavior could be due to the fluorescence at low concentration being below the threshold for which the machine can measure fluorescence reliably or the dye preferentially binding to the walls of each well. It could also be due to spatial differences associated with the top row in the 96-well plate (see Table S1 in the SM) where the low concentration wells are positioned.

To determine whether the behavior at low dye concentrations was due to the position of the wells or the low signal, an additional experiment was performed with well concentrations rotated so that the dye concentrations increased from right-to-left and from bottom-to-top (see Table S2 in the SM). In this experiment, the same behavior was observed at low dye concentration (see Figure S1 in the SM). As a result, we attribute the behavior at low dye concentration to the signal being below the range of the instrument. The minimum fluorescence value that can be measured reliably, or *F*_min_, was determined by calculating the largest fluorescence value observed at the lowest temperature for all data points associated with dye concentrations below 0.5 *μ*mol/L. This results in *F*_min_ = 0.23 *×* 10^6^.

The fluorescence trends of double-stranded DNA are slightly different than those of single-stranded DNA. At dye concentrations above 2.5 *μ*mol/L, the fluorescence becomes a nonlinear function of dye concentration. This concentration of 2.5 *μ*mol/L was also noted to be the approximate upper limit of linearity with respect to SYBR Gold (7). As discussed in previously, nonlinearities in fluorescence could be due to quenching, non-dilute adsorption, deviation from Beer’s Law, or any combination of the three. To avoid such effects, which are not accounted for in the model, data points for DS with dye concentrations above 2.5 *μ*mol/L were excluded. Having selected each dye concentration regime as described above for each DNA concentration *k* and type *t*, the concentrations outside the regime were removed from the indices of 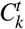 and the columns of 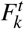.

At very high temperature, the fluorescence signal becomes very small, eventually decreasing below *F*_min_. To identify temperatures that have fluorescence values that are too small to be reliable, we determined the largest temperature *T*_max_ such that the average fluorescence over all wells of all plates was above *F*_min_. This resulted in *T*_max_ being 316.5 K, 324.5 K, 322 K, and 329 K for SS with *D* = 1.0 *μ*mol/L, SS with *D* = 2.0 *μ*mol/L, DS with *D* = 1.0 *μ*mol/L, and DS with *D* = 2.0 *μ*mol/L, respectively. Data points above this temperature were excluded from further investigation by removing their row in 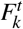. Calculations from the NUPACK web application(20) indicate that no double-stranded DNA melts in these temperature regimes.

Before using (8) to calculate thermodynamic and photophysical properties with numerical optimization, the dimensionless variables

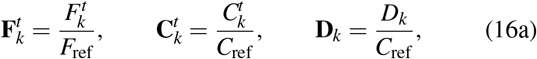

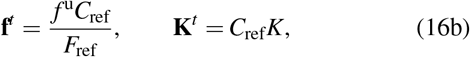

were introduced for each DNA concentration index *k* = 1, 2. Here, 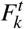 is an *m × n* matrix containing the *m* temperatures and *n* wells in the regime determined above for each *k* and *t*, while 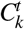 is a column vector containing the *n* wells. The quantities **f**^*t*^ and **K**^*t*^ are column vectors containing the *m* temperatures, and **D** is a scalar. In this work, we set *F*_ref_ = 1 *×* 10^6^ and *C*_ref_ = 1 *μ*mol/L. Since this choice makes **D** = *k*, we will use **D** instead of *k* in subscripts, and drop the subscript on **D**. Since we will investigate SS and DS independently, we will drop the superscript *t*.

In vectorized form and dimensionless quantities with this notation, (8) becomes

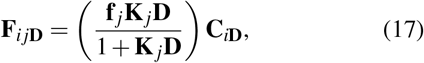

for each well *i* = 1 to *n*, each temperature index *j* = 1 to *m*, and each DNA concentration **D** = 1 or 2. In writing (17), the assumption of negligible molar fluorescence of dye in solution, or (15), has been used.

### Noise Removal

As the raw data exhibited significant well-to-well and plateto-plate variation, it is useful to remove the noise before calculating the parameters. Since the experimental data is expected to be linear in dye concentration at constant DNA concentration and temperature, one approach is to recast the noisy data as its closest linear approximation. Using **M**_**D**_ to denote the column vector of slopes at a DNA concentration **D** for each temperature, we seek to find an approximation

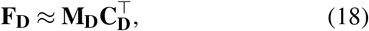

where **M**_**D**_ = (**M**_1**D**_, …, **M**_*m***D**_)^T^ and

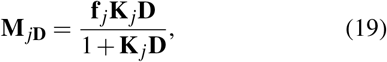

for each temperature *T*_*j*_ and DNA concentration **D**. Since the approach for calculation of **M**_**D**_ will be independent of **D**, we will drop the subscript in this section to simplify notation.

A conventional approach is to postulate that the experimental data is not linear due to error in the fluorescence signal, and to calculate **M** so as to minimize error between the observed signal 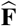 and the theoretical signal **F**. This is a Least Squares (LS) regression problem which may be expressed as

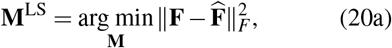

subject to

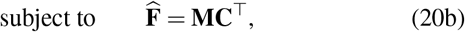

where || · ||_*F*_ is the Frobenius norm and the decision variable **M** is a column vector with *m* entries. The analytical solution to (20) is

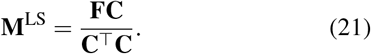

In order to determine whether **M**^LS^ is realistic, it is important to assess the validity of the approximation (18) by assessing the error between **F** and 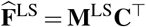. In Figure 3, 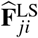 is plotted as a function of **F**_*ji*_ for each temperature *T*_*j*_ and concentration **C**_*i*_. The subplots are organized in the same manner as Figure 2. The trajectories along which the temperature appears to change linearly are interpreted to be associated with a single well.

In general, the difference between **F** and 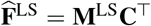 is significant and unphysical. Several replicates differ in the sign of the sum of residuals 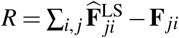. For example, replicate (SS,A,1) has *R* = *−* 1058. This means that, on the day that the experiment was run, the machine was systematically *underestimating* the signal by a significant magnitude. Replicate (SS,B,1), however, has *R* = 739. This implies that, on the different day in which the latter experiment was run, the machine was systematically *overestimating*. Taken together, these trends are unrealistic. A machine that is routinely calibrated should have a relatively small error in signal that is nearly the same from one plate to the next. The paths of temperature gradient for each well suggest that an error specific to each well has not been considered in the least squares approach.

This source of noise could arise from imperfect transferring of liquid volumes. As a result, the total concentration of DNA or dye in each well could be different than the theoretically set value. Since all wells in each plate were given the same DNA concentration, but different dye concentrations, we hypothesize that the error associated with each well is due primarily to an error in dye concentration. Unlike the fluorescence signal, each dye concentration is not measured directly in each well.

Postulating that the experimental data is not linear due to error *both* in **F** and **C**, we seek to minimize the sum of the errors in observed signals and concentrations as

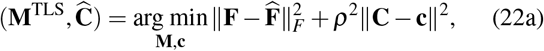

subject to

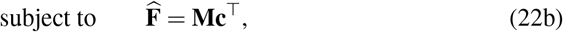

where *ρ* is a parameter representing the weight attributed to the error in concentration, || · ||is the Euclidian norm, **M** is a column vector of length *m*, and **c** is a column vector of length *n*. In contrast to (20b), where 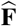 is linear in the decision variables, 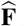 in Equation (22b) is bilinear. This optimization problem is analogous to what has been described as Total Least Squares (TLS) (see, e.g., Ref. (21)). The quantities **M**^TLS^ and **Ĉ**, as well as their variances, *V* (**M**) and *V* (**Ĉ**), respectively, are calculated as described in Section S1.2 of the SM.

The error in fluorescence signal associated with the total least squares model is depicted in Figure 4. As expected, the error in signal is quite small and does not appear to differ greatly from one replicate measurement to the next. The largest deviations appear to occur at low temperature when the signal is large (see, for example, Figure 4(SS,A,2) and Figure 4(DS,B,2)). In these regimes, the model systematically overestimates the fluorescence signal.

**Figure 4:**
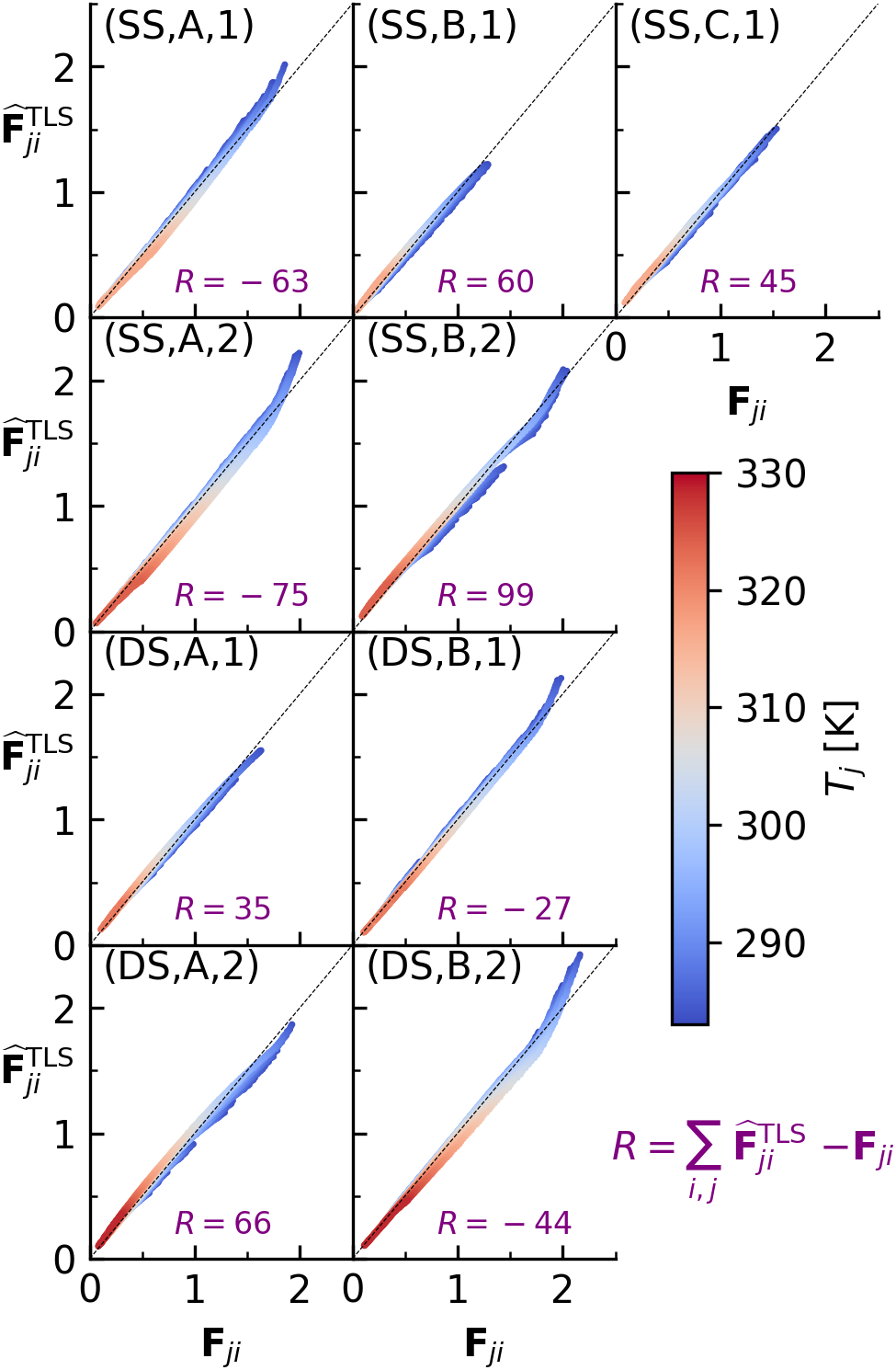
As Figure 3, with 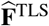 instead of 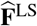.

It is somewhat expected that the model would systematically overestimate the signal only when the signal is large. When the signal is large, the dye concentration is large and/or the temperature is small. In this regime, quenching, deviations from Beer’s Law, as well as non-dilute dye adsorption may occur. When any of these phenomena occur, the model will overestimate the signal.

The differences between the total dye concentrations calculated from mass balances representing dilutions, **C**, and from total least squares, **Ĉ**, are depicted in Figure 5. In contrast to the previous figures, results from the same replicates are depicted in the same subplot.

**Figure 5:**
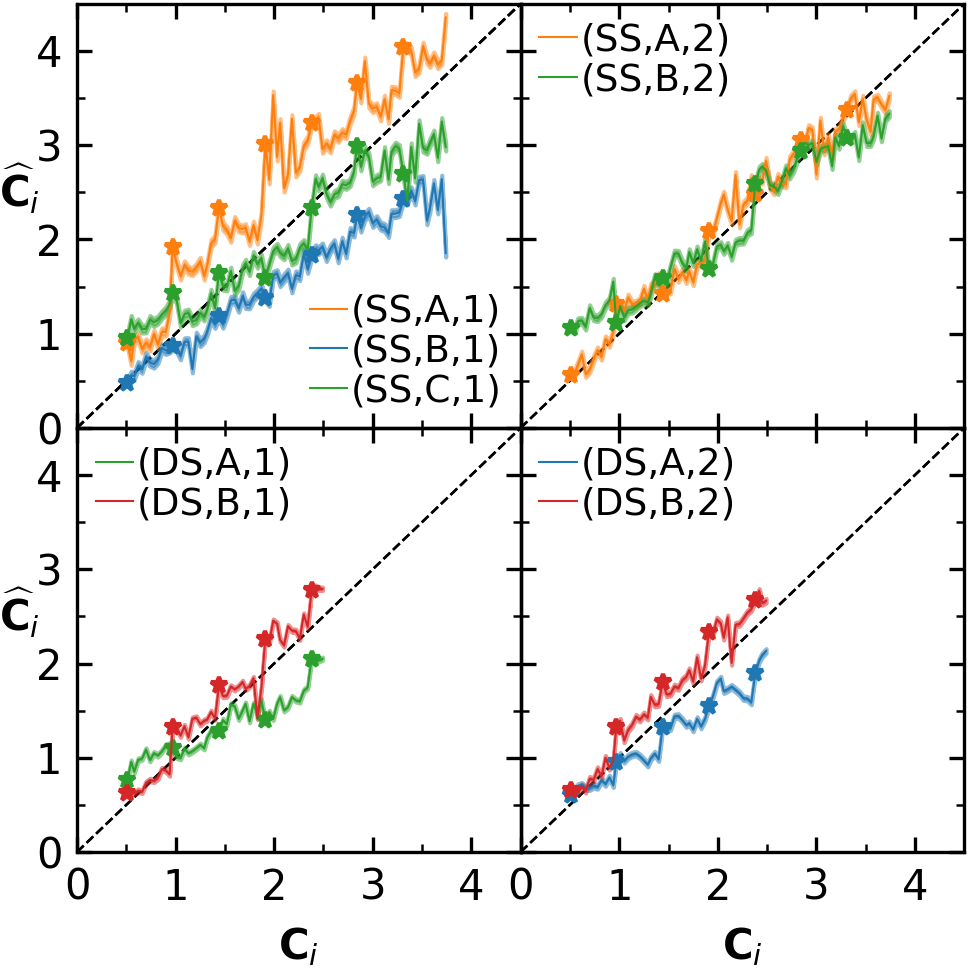
Concentration calculated from total least squares, **Ĉ**_*i*_, as a function of signal calculated experimentally, or **C**_*i*_, for each well *i*. The colors correspond to different replicate measurements, labelled as in Table 1. The points marked with a star (*\**) correspond to column 1 in the 96-well plate. Each color (i.e., red, green, orange, or blue) groups a set of plates that used the same working concentrations. The shadings correspond to 3 standard deviations about **Ĉ**_*i*_.

The differences between the concentrations from dilution balances and from total least squares are consistent with the pipetting protocol. In many cases, the average difference between **Ĉ**_*i*_ and **C**_*i*_ is highly correlated with replicate (i.e., color in each subplot). This is attributed to each replicate using the same set of working concentrations (see Section S1.1 in SM). The colors in Figure 5 correspond to plates whose dye concentrations were also prepared from identical sets of working concentrations. In most cases, the results from different plates that were prepared using the same working concentrations appear to be quite correlated. Finally, the noisy variations for data points between each *\** of the same dataset in Figure 5 correspond to a row in a 96-well plate. This is also consistent with the pipetting procedure (see Section S1.1 in the SM), since the differences in concentration along a row were generated from a working solution prepared from an extra serial dilution.

The fluorescence–concentration slopes, calculated by total least squares, are depicted in Figure 6 for each temperature and DNA concentration. For each type of DNA, the slope increases with increasing DNA concentration. In all cases, 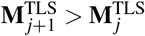. That is, the slopes increase with decreasing temperature. The largest estimated error in **M**^TLS^ occurs at low temperature where the fluorescence value is highest. This error is attributed to a lack of accounting for nonlinearities in the model, which results in the model overestimating the fluorescence signal (see also Figure 4). Since the slope **M** increases with increasing DNA concentration, the theoretical signal 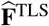 also increases with increasing DNA concentration at fixed dye concentration. As more DNA is added into solution, more dye partitions from solution into DNA, and the fluorescence intensity is increased.

**Figure 6:**
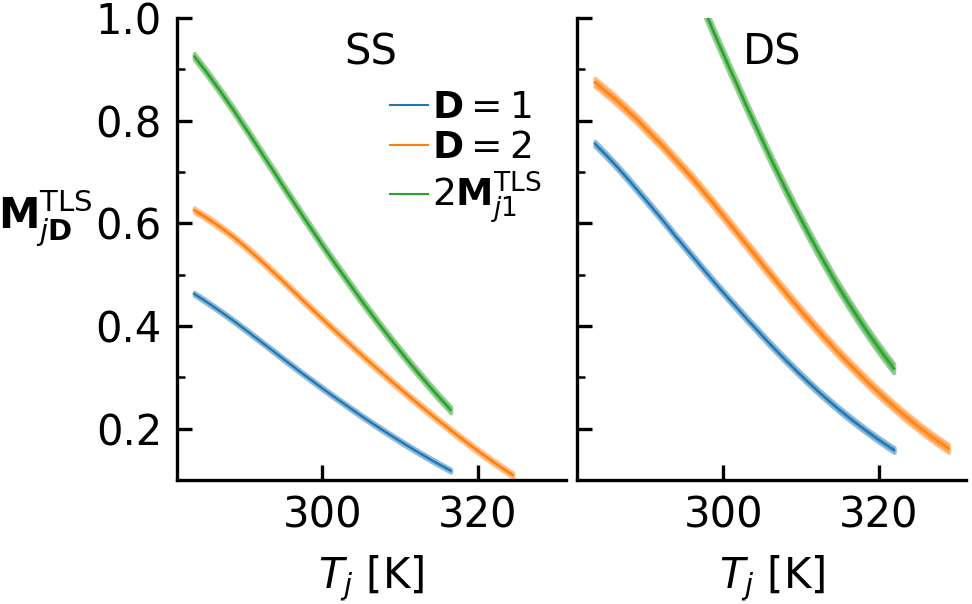
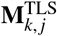 for each temperature *T*_*j*_ associated with SS (left) and DS (right) at DNA concentrations of **D** = 1 (blue) and **D** = 2 (orange). The shaded regions at each temperature *T*_*j*_ depict 3 standard deviations about 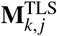. The green lines depict twice the value obtained at the lower concentration of DNA.

### Binding Strength and Molar Fluorescence

Having removed the noise in the experimental data, the second step in our procedure of calculating thermostatic and photophysical properties from fluorescence measurements involves calculating the binding strength **K** and molar fluorescence **f** from **M**_**D**_. The challenge is that both **K** and **f** depend on temperature, and both decrease with temperature. In Equation (19), it is challenging to deconvolute **K** and **f** using **M**_**D**_ alone.

One option is to exploit the dependence of **M**_**D**_ on DNA concentration **D**. Rearranging (19) leads to

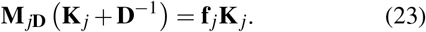

Here, the right-hand-side is independent of **D**. Having calculated **M**_**D**_ for different **D**, as was done in the previous section (omitting the superscript TLS for brevity), **K** may be calculated by minimizing the difference of the left-hand-side calculated at different **D**. In general, the procedure described in this section can be extended quite readily to any finite number of DNA concentrations. For simplicity, however, we only present the description for **D** = 1 and 2, which were considered in this work. This corresponds to the following optimization problem

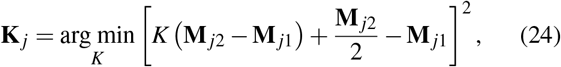

for each *j* = 1 to *m*, where *K* is a scalar. Since more temperature indices on **M**_2_ were originally obtained than **M**_1_ (see horizontal axis range in Figure 6), **M**_2_ has been redefined as a subset of itself containing only the indices associated with **M**_1_. Equation (24) is a linear least squares problem. Since **M** _*j*2_ ≠ **M** _*j*1_ for each temperature *T*_*j*_ (see Figure 6), the objective function is a convex quadratic. Its unique solution is

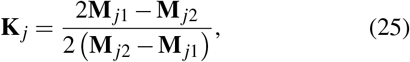

for each temperature *T*_*j*_. Since **M** _*j*2_ *>* **M** _*j*1_, and 2**M** _*j*1_ *>* **M** _*j*2_ (see Figure 6), Equation (25) demonstrates that **K** _*j*_ *>* 0. In other words, the partition coefficient is always positive.

Having calculated **K** via (25), **f** may be calculated from substituting the value into (23) and rearranging. However, the value calculated depends on the DNA concentration chosen (i.e., **D** = 1 or 2). For this overdetermined system, *f* may be calculated in a least squares sense via

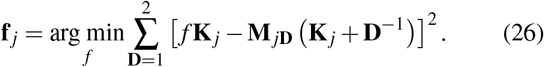

for each temperature *T*_*j*_, where *f* is a scalar. Equation (25) is also a linear least squares problem. Since **K** _*j*_ ≠ 0, the objective function is also a convex quadratic that admits the unique solution

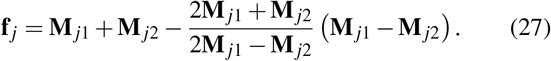

The results of computations from (25) and (27) are depicted in Figure 7. The partition coefficients decrease with increasing temperature. This is expected for adsorption, which is accompanied by a release of heat (exothermic). The adsorbed fluorescences also decrease with increasing temperature (except for SS at the second lowest temperature investigated, where a small increase *<* 10^*−*4^ is observed). While the relationship between the molar fluorescence of a single compound and temperature is complex, the decrease with temperature is characteristic of a brightness that decreases with temperature.

**Figure 7:**
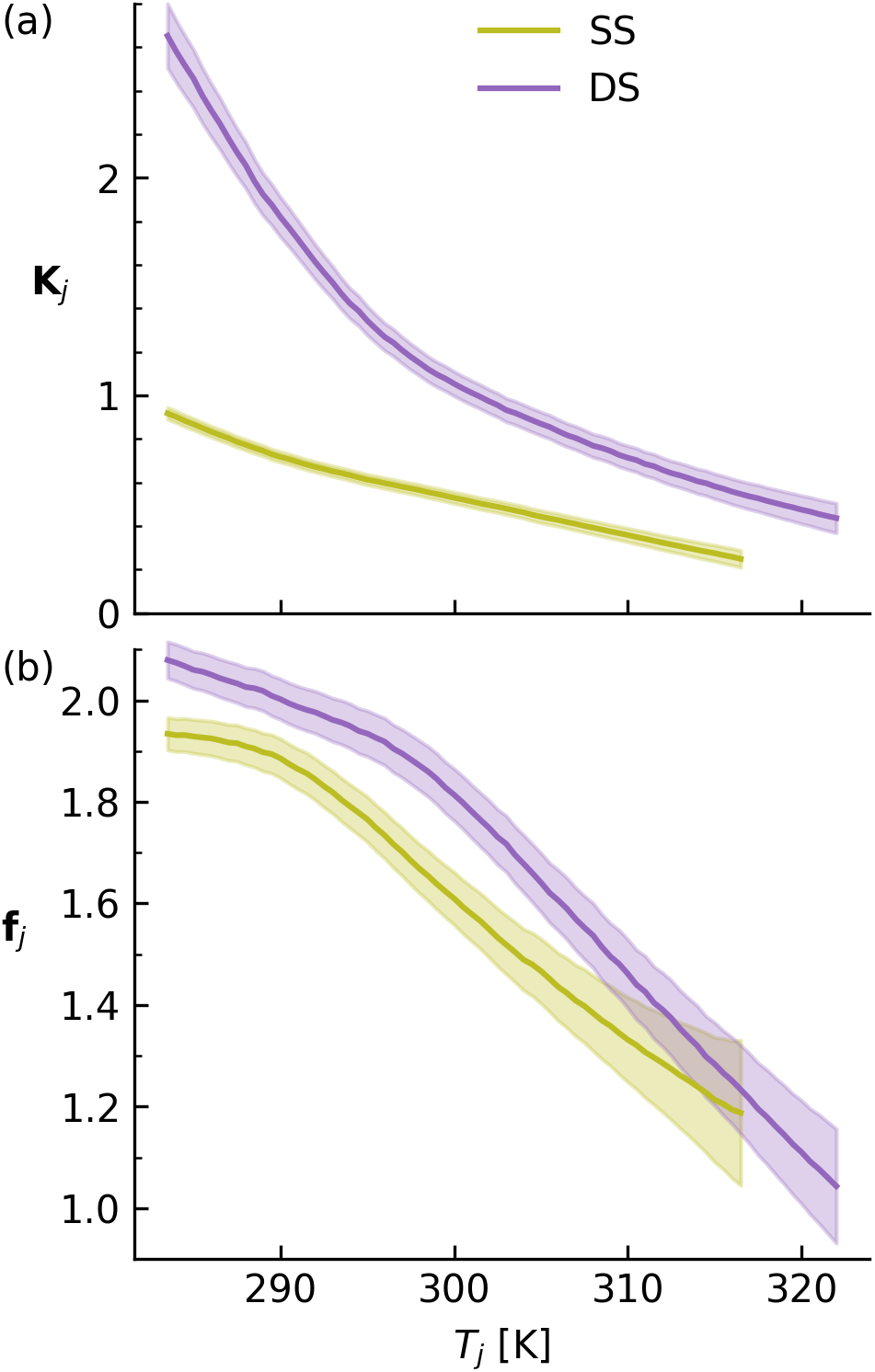
(a) Partition coefficient **K**_*j*_ calculated from (25) and (b) adsorbed fluorescence **f**_*j*_ calculated from (27) at each temperature *T*_*j*_ for SS (yellow) and DS (purple). The shaded regions correspond to 3 standard deviations about the mean, estimated with *V* (**M**) and a linear approximation for division in (25) or (27).

The calculations of **f** and **K** both for SS and DS were performed independently of one another. Yet, the results are consistent with biochemical intuition. Adsorption of dye onto

DS is stronger than SS, and fluorescence of dye adsorbed to DS is higher than SS. The larger partition coefficient for DS corresponds to a larger fraction of bound dye of *≈* 0.2 (see Figure S3 in the SM). Except at very large dye concentrations and very low temperatures, the average number of dye bound to each DNA strand is less than 2 (see Figure S4 in the SM). As the number of base pairs per DNA strand is 22, this validates the assumption that the number of free adsorption sites is constant.

Deconvoluting binding strength from molar fluorescence explains why the total fluorescence of DS is larger than that of SS. The explanation, in fact, depends on the temperature of interest. At temperatures above 305 K, the binding strength onto DS is still larger than SS; however, the molar fluorescence has no statistically significant difference. In contrast to the low temperature regime, the difference in total fluorescence at high temperature is driven only by differences in binding. While the fluorescence per dye adsorbed is the same, the number of dye adsorbed is different.

Examination of (2) demonstrates that another useful property can be obtained. From (2), the quotient of the molar fluorescence in one phase to another is the quotient of the brightness. Specifically,

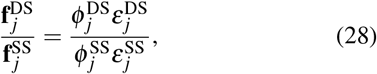

for each temperature index *j* available for SS, where the superscripts SS and DS have been reintroduced. The results, depicted in Figure S5 in the SM, demonstrate that, depending on the temperature, the dye adsorbed to DS is usually approximately 5 to 10 % more bright. In practice, this approach of calculating the relative brightness may be applied to any ligand–receptor combination with negligible spectral overlap.

As compared to SYBR Gold and SYBR Green I, the adsorption strength of SYTO-13 onto DS is smaller (see Table 2) by a factor of about one order-of-magnitude. This is consistent with a previous report(18) which stated that, in contrast to SYBR Green I, SYTO-13 does not inhibit PCR and does not influence melting temperature. However, since PCR is a temperature-dependent process, the inhibition of PCR due to dye binding is more thoroughly characterized with the temperature dependence of the association constant. Determining the temperature-dependence of the binding of SYBR Gold and SYBR Green I is therefore a useful direction for future work.

**Table 1:** Abbreviations Used for Data Sets

| $t^a$ | $\ell^b$ | $k^c$ | Name |
| --- | --- | --- | --- |
| SS | A | 1 | (SS,A,1) |
| SS | B | 1 | (SS,B,1) |
| SS | C | 1 | (SS,C,1) |
| SS | A | 2 | (SS,A,2) |
| SS | B | 2 | (SS,B,2) |
| DS | A | 1 | (DS,A,1) |
| DS | B | 1 | (DS,B,1) |
| DS | A | 2 | (DS,A,2) |
| DS | B | 2 | (DS,B,2) |
| None | A | 0 | (A,0) |

<sup>a</sup> Type of DNA.
<sup>b</sup> Name of Replicate.
<sup>c</sup> DNA concentration index, $\lfloor D \rfloor$ .

**Table 2:** Association Constants of Dyes at Room Temperature or 295 K

| Name of Dye | $K_a = \frac{K}{2} \left[ \frac{\mu\text{mol}}{\text{L}} \right]$ | Reference |
| --- | --- | --- |
| SYBR Gold<sup>a</sup> | 3.66 | Ref. (7) |
| SYBR Gold<sup>b</sup> | 7.19 | Ref. (7) |
| SYBR Gold<sup>b</sup> | 6.45 | Ref. (7) |
| SYBR Gold<sup>a,b</sup> | $7.8 \pm 3.3$ | Ref. (22) |
| SYBR Green I<sup>a</sup> | 3.00 | Ref. (7) |
| SYBR Green I<sup>b</sup> | 22.2 | Ref. (3) |
| SYTO-13<sup>b</sup> | $0.67 \pm 0.03$ | This work |

<sup>a</sup> Magnetic tweezers force extension
<sup>b</sup> Fluorescence

### Binding Thermodynamics

By obtaining the temperature-dependence of the partition coefficients and the molar fluorescences, an additional third step may be undertaken to compute enthalpies of binding and relative brightness. The free energy of adsorption at each temperature index *j* may be calculated as

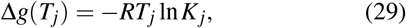

where *R* is the gas constant. For an ideal solution with a small coefficient of thermal expansion and equilibrium adsorption dictated by (6), the enthalpy of adsorption Δ*h* is

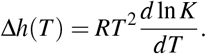

However, since *T* and *P* are only known at discrete values, an approximation is needed to compute Δ*h*_*j*_ = Δ*h*(*T*_*j*_) for each *j*. Using a central-finite difference approximation for temperatures equally spaced by Δ*T*, one obtains

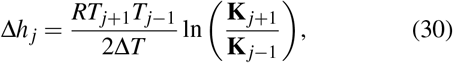

for *j* = 2, …, *m−* 1. From (29) and (30), the entropy of dye adsorption may be calculated from Δ*s*_*j*_ = (Δ*h*_*j*_ *−* Δ*g*_*j*_) */T*_*j*_ for *j* = 2, …, *m−* 1.

The free energies and enthalpies of adsorption are depicted in Figure 8. For each type of DNA, the free energy of binding is nearly independent of temperature. The free energy of binding of SYTO-13 at 295 K to SS is *−* 32.68 *±* 0.09 kJ/mol, while that to DS is *−* 34.60 *±* 0.11 kJ/mol. The smaller Δ*g* for DS reflects the larger partition coefficient.

**Figure 8:**
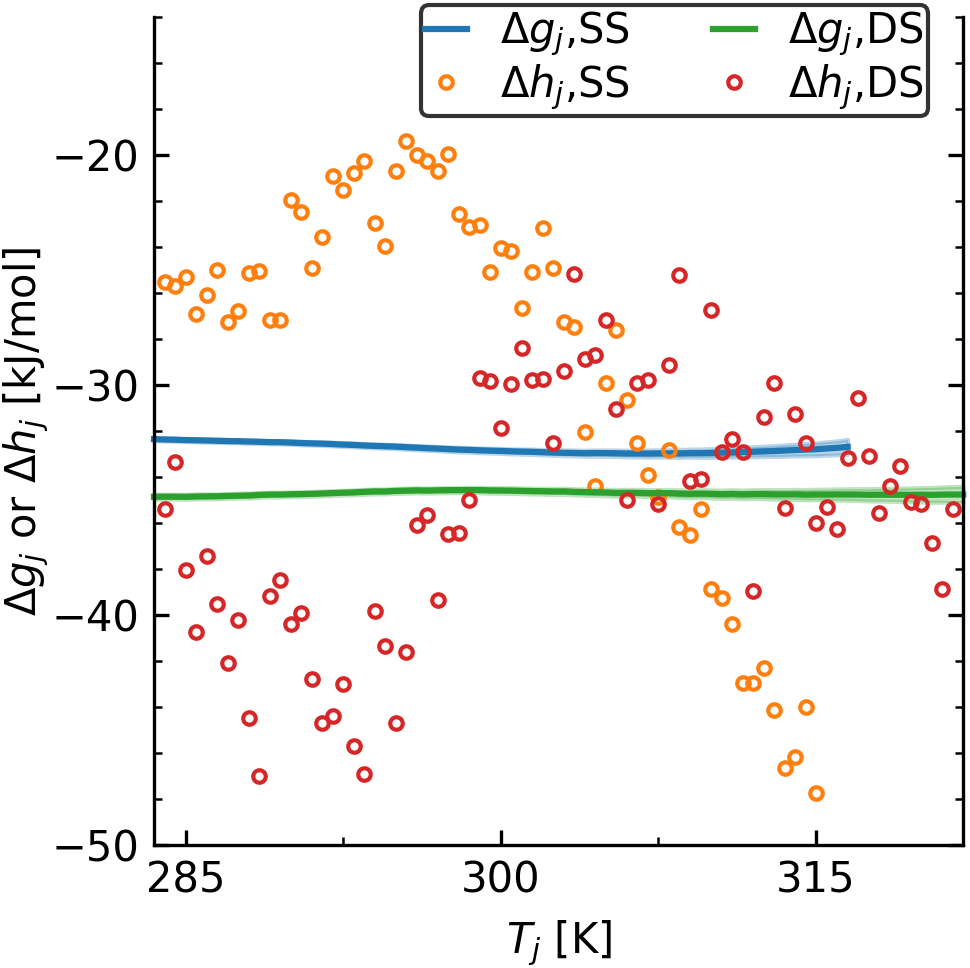
Free energy of transfer (lines, Equation (29)) or enthalpy of transfer (circles, Equation (30)) for dye from solution to SS or DS as a function of temperature.

The enthalpy of binding is calculated less accurately than the free energy of binding, as (30) is a finite-difference approximation. The enthalpy of binding of dye to DS, as depicted in circles in the right subplot of Figure 8, appears to waver above and below an average value along temperature. This means that the heat capacity difference of dye between both phases,

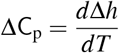

is small. The mean value of Δ*h*_*j*_ for dye binding to DS over all temperatures is *−* 35 *±* 5 kJ/mol. The enthalpy of binding to SS also appears to waver about a mean value at temperatures below 300 K. However, at high temperature, the value calculated from (30) continues to decrease. This corresponds to a negative heat capacity of adsorption, or ΔC_p_ *<* 0.

## CONCLUSION

In this work, a protocol for extracting temperature-dependent thermodynamic and photophysical properties of fluorescent dyes interacting with DNA was presented. The fluorescence was measured at a large number of state points by using a PCR thermocycler, while properties were extracted in a robust and efficient manner with numerical optimization. While applied to SYTO-13 interacting with a 22 base-pair oligonucleotide, the technique can be readily extended to any ligand–receptor combination possessing negligible spectral overlap. To facilitate reproducibility of the approach, as well as its transferability and further extension, we have made the code available at https://github.com/usnistgov/dye_dna_plates.

A key aspect of the approach involves exploiting the linear relationship between fluorescence and dye concentration that occurs at dilute concentrations. In this linear regime, it is guaranteed that dye quenching and dimerization are negligible. While the approach is not able to capture cooperative binding of dye to DNA, such behavior is usually not important. The end goal of many assays is to quantify concentrations of DNA in solution; the dye is employed only as a reporter to achieve this goal. Taking advantage of the dilute regime can reduce bias in quantification.

Another distinguishing aspect of the technique is that it does not require experimental data to look pretty; that is, it can account for data that exhibits significant well-to-well and plate-to-plate variation. This provides the technique with increased throughput. Rather than discarding data possessing noisy oscillations, the approach can relate the trends to specific experimental protocol.

In contrast to previous studies, the procedure emphasized the utility of extracting the molar fluorescence. This photophysical property is necessary for quantification. While its value is instrument-dependent, it can be used to determine the brightness relative to a known standard. At the same wavelengths, the brightness is transferable between instruments. In order to explain why the fluorescence of a certain dye–DNA solution is larger than another dye–DNA solution, it is necessary to discriminate between molar fluorescence and binding strength.

Temperature-dependent thermodynamic and photophysical properties, as can be produced by the technique reported in this work, are useful. Properties like the enthalpy of binding provide helpful intuition on the nature of dye–DNA interactions. In addition, such properties can enable quantification to be less empirical. Assumptions made in the absence of available data can be verified or relaxed entirely.

## AUTHOR CONTRIBUTIONS

All authors contributed to design of experiments and conceptualization of the project. R. F. D. developed mathematical model. J. A. L. performed experiments. R. F. D. and A. J. K. established method for data analysis, and R. F. D. wrote and tested all computer code. R. F. D., J. A. L, and A. J. K analyzed the results of calculations. All authors contributed to preparation of manuscript.

## ACKNOWLEDGMENTS

R. F. D. acknowledges support from a National Research Council fellowship and the NIST Professional Research Experience Program through University of Maryland. R. F. D. thanks R. Luke for careful reading and suggestions associated with an earlier draft of this work.

## SUPPORTING MATERIAL

An online supplement to this article can be found by visiting BJ Online at http://www.biophysj.org.

## Footnotes

1 Certain commercial equipment, instruments, software, or materials are identified in this paper in order to specify the experimental procedure adequately. Such identification is not intended to imply recommendation or endorsement by the National Institute of Standards and Technology, nor is it intended to imply that the materials or equipment identified are necessarily the best available for the purpose.

## REFERENCES

1. Ihmels, H., and D. Otto, 2005. Intercalation of Organic Dye Molecules into Double-Stranded DNA General Principles and Recent Developments, Springer-Verlag Berlin, volume 258, 161–204.

2. Biver, T., A. De Biasi, F. Secco, M. Venturini, and S. Yarmoluk, 2005. Cyanine Dyes as Intercalating Agents: Kinetic and Thermodynamic Studies on the DNA/Cyan40 and DNA/CCyan2 Systems. Biophys. J. 89:374–383.

3. Dragan, A. I., J. R. Casas-Finet, E. S. Bishop, R. J. Strouse, M. A. Schenerman, and C. D. Geddes, 2010. Characterization of PicoGreen Interaction with dsDNA and the Origin of Its Fluorescence Enhancement upon Binding. Biophys. J. 99:3010–3019.

4. Dragan, A. I., R. Pavlovic, J. B. McGivney, J. R. Casas-Finet, E. S. Bishop, R. J. Strouse, M. A. Schenerman, and C. D. Geddes, 2012. SYBR Green I: Fluorescence Properties and Interaction with DNA. J. Fluoresc. 22:1189–1199.

5. Islam, M. M., M. Chakraborty, P. Pandya, A. Al Masum, N. Gupta, and S. Mukhopadhyay, 2013. Binding of DNA with Rhodamine B: Spectroscopic and Molecular Modeling Studies. Dyes Pigm. 99:412–422.

6. Paul, P., and G. S. Kumar, 2013. Spectroscopic Studies on the Binding Interaction of Phenothiazinium Dyes Toluidine Blue O, Azure A and Azure B to DNA. Spectrochim. Acta A Mol. Biomol. Spectrosc. 107:303–310.

7. Kolbeck, P. J., W. Vanderlinden, G. Gemmecker, C. Gebhardt, M. Lehmann, A. Lak, T. Nicolaus, T. Cordes, and J. Lipfert, 2021. Molecular Structure, DNA Binding Mode, Photophysical Properties and Recommendations for Use of SYBR Gold. Nucleic Acids Res. 49:5143–5158.

8. Gaugain, B., J. Barbet, N. Capelle, B. P. Roques, and J.-B. Le Pecq, 1978. DNA Bifunctional Intercalators. 2. Fluorescence Properties and DNA Binding Interaction of an Ethidium Homodimer and an Acridine Ethidium Heterodimer. Biochem. 17:5078–5088.

9. Georges, J., 1995. Deviations from Beer’s Law Due to Dimerization Equilibria: Theoretical Comparison of Absorbance, Fluorescence and Thermal Lens Measurements. Spectrochim. Acta 51A:985–995.

10. Le Pecq, J.-B., M. Le Bret, J. Barbet, and B. Roques, 1975. DNA Polyintercalating Drugs: DNA Binding of Diacridine Derivatives. Proc. Nat. Acad. Sci. USA 72:2915–2919.

11. Choi, Y.-J., and K. Sawada, 2021. Fluorescence Sensors. In Reference Module in Biomedical Sciences, Elsevier.

12. Levitus, M., 2020. Tutorial: Measurement of Fluorescence Spectra and Determination of Relative Fluorescence Quantum Yields of Transparent Samples. Methods Appl. Fluoresc. 8.

13. McGhee, J. D., and P. H. von Hippel, 1974. Theoretical Aspects of DNA-Protein Interactions: Co-operative and Non-co-operative Binding of Large Ligands to a Onedimensional Homogeneous Lattice. J. Mol. Biol. 86:469–489.

14. Villaluenga, J. P. G., J. Vidal, and F. J. Cao-García, 2020. Noncooperative Thermodynamics and Kinetic Models of Ligand Binding to Polymers: Connecting McGhee–von Hippel Model with the Tonks Gas Model. Phys. Rev. E 102:012407.

15. Benesi, H. A., and J. H. Hildebrand, 1949. A Spectrophotometric Investigation of the Interaction of Iodine with Aromatic Hydrocarbons. J. Am. Chem. Soc. 71:2703–2707.

16. Peacocke, A. R., and J. N. H. Skerrett, 1955. The Interaction of Aminoacridines with Nucleic Acids. Trans. Faraday Soc. 52:261–279.

17. Palais, R., and C. T. Wittwer, 2009. Chapter 13 Mathematical Algorithms for High-Resolution DNA Melting Analysis. Methods Enzymol. 454:323–343.

18. Gudnason, H., M. Dufva, D. D. Band, and A. Wolff, 2007. Comparison of Multiple DNA Dyes for Real-Time PCR: Effects of Dye Concentration and Sequence Composition on DNA Amplification and Melting Temperature. Nucleic Acids Res. 35:e127.

19. Radvanszky, J., M. Surovy, E. Nagyova, G. Minarik, and L. Kadasi, 2015. Comparison of Different DNA Binding Fluorescent Dyes for Applications of High-Resolution Melting Analysis. Clin. Biochem. 48:609–616.

20. Zadeh, J. N., C. D. Steenberg, J. S. Bois, B. R. Wolfe, M. B. Pierce, A. R. Khan, R. M. Dirks, and N. A. Pierce, 2011. NUPACK: Analysis and Design of Nucleic Acid Systems. J. Comput. Chem. 32:170–173.

21. Golub, G. H., and C. F. van Loan, 1980. An Analysis of the Total Least Squares Problem. SIAM J. Numer. Anal. 17:883–893.

22. Biebricher, A. S., I. Heler, R. F. H. Roijmans, T. P. Hoekstra, E. J. G. Peterman, and G. J. L. Wuite, 2015. The Impact of DNA Intercalators on DNA and DNA-Processing Enzymes Elucidated Through Force-Dependent Binding Kinetics. Nat. Commun. 6:7304.

