## Supplementary Material for "Temperature-dependent Thermodynamic and Photophysical Properties of SYTO-13 Dye Bound to DNA"

### Contents

**S1 Supplemental Methods**

S1.1 Experimentation . . . . . . . . . . . . . . . . . . . . . . . . . . . . . . . . . . . . . . . . .

S1.2 Total Least Squares . . . . . . . . . . . . . . . . . . . . . . . . . . . . . . . . . . . . . . .

**S2 Supplemental Results**

S2.1 Fraction of Bound Dye . . . . . . . . . . . . . . . . . . . . . . . . . . . . . . . . . . . . . .

S2.2 Average Dye per DNA . . . . . . . . . . . . . . . . . . . . . . . . . . . . . . . . . . . . . .

### S1 Supplemental Methods

#### S1.1 Experimentation

Table S1: Concentrations of dye ($\mu$mol/L) used for each well in 96-well plate.

| | 1 | 2 | 3 | 4 | 5 | 6 | 7 | 8 | 9 | 10 | 11 | 12 |
|---|---|---|---|---|---|---|---|---|---|---|---|---|
| A | 0.04 | 0.08 | 0.12 | 0.16 | 0.19 | 0.23 | 0.27 | 0.31 | 0.35 | 0.39 | 0.43 | 0.47 |
| B | 0.51 | 0.55 | 0.58 | 0.62 | 0.66 | 0.70 | 0.74 | 0.78 | 0.82 | 0.86 | 0.90 | 0.94 |
| C | 0.97 | 1.01 | 1.05 | 1.09 | 1.13 | 1.17 | 1.21 | 1.25 | 1.29 | 1.32 | 1.36 | 1.40 |
| D | 1.44 | 1.48 | 1.52 | 1.56 | 1.60 | 1.64 | 1.68 | 1.71 | 1.75 | 1.79 | 1.83 | 1.87 |
| E | 1.91 | 1.95 | 1.99 | 2.03 | 2.06 | 2.10 | 2.14 | 2.18 | 2.22 | 2.26 | 2.30 | 2.34 |
| F | 2.38 | 2.42 | 2.45 | 2.49 | 2.53 | 2.57 | 2.61 | 2.65 | 2.69 | 2.73 | 2.77 | 2.81 |
| G | 2.84 | 2.88 | 2.92 | 2.96 | 3.00 | 3.04 | 3.08 | 3.12 | 3.16 | 3.19 | 3.23 | 3.27 |
| H | 3.31 | 3.35 | 3.39 | 3.43 | 3.47 | 3.51 | 3.55 | 3.58 | 3.62 | 3.66 | 3.70 | 3.74 |

The spatial distribution of dye concentrations, an 8 by 12 matrix $X$, in each plate is depicted in Table S1. It was prepared via

$$X = v(A + B),$$

where $A$ is an $8 \times 12$ matrix containing

$$A = \begin{bmatrix} a & \dots & a \end{bmatrix}, \qquad a = (0, 1.10, 2.20, 3.30, 4.40, 5.50, 6.60, 7.70)^\top,$$

$B$ is an $8 \times 12$ matrix containing

$$B = \begin{bmatrix} b \\ \vdots \\ b \end{bmatrix}, \qquad b = (0.09, 0.18, 0.28, 0.37, 0.46, 0.55, 0.64, 0.73, 0.83, 0.92, 1.01, 1.10),$$

and $v = 0.425$ is the final volume fraction in each well. The working column $a$ of solutions with concentrations in $\mu$mol/L was generated by dilution from stock, while the working row $b$ was prepared after one additional dilution step.

Table S2: Concentrations of dye ($\mu$mol/L) used for each well in rotated plate.

| | 1 | 2 | 3 | 4 | 5 | 6 | 7 | 8 | 9 | 10 | 11 | 12 |
|---|---|---|---|---|---|---|---|---|---|---|---|---|
| A | 3.74 | 3.70 | 3.66 | 3.62 | 3.58 | 3.55 | 3.51 | 3.47 | 3.43 | 3.39 | 3.35 | 3.31 |
| B | 3.27 | 3.23 | 3.19 | 3.16 | 3.12 | 3.08 | 3.04 | 3.00 | 2.96 | 2.92 | 2.88 | 2.84 |
| C | 2.81 | 2.77 | 2.73 | 2.69 | 2.65 | 2.61 | 2.57 | 2.53 | 2.49 | 2.45 | 2.42 | 2.38 |
| D | 2.34 | 2.30 | 2.26 | 2.22 | 2.18 | 2.14 | 2.10 | 2.06 | 2.03 | 1.99 | 1.95 | 1.91 |
| E | 1.87 | 1.83 | 1.79 | 1.75 | 1.71 | 1.68 | 1.64 | 1.60 | 1.56 | 1.52 | 1.48 | 1.44 |
| F | 1.40 | 1.36 | 1.32 | 1.29 | 1.25 | 1.21 | 1.17 | 1.13 | 1.09 | 1.05 | 1.01 | 0.97 |
| G | 0.94 | 0.90 | 0.86 | 0.82 | 0.78 | 0.74 | 0.70 | 0.66 | 0.62 | 0.58 | 0.55 | 0.51 |
| H | 0.47 | 0.43 | 0.39 | 0.35 | 0.31 | 0.27 | 0.23 | 0.19 | 0.16 | 0.12 | 0.08 | 0.04 |

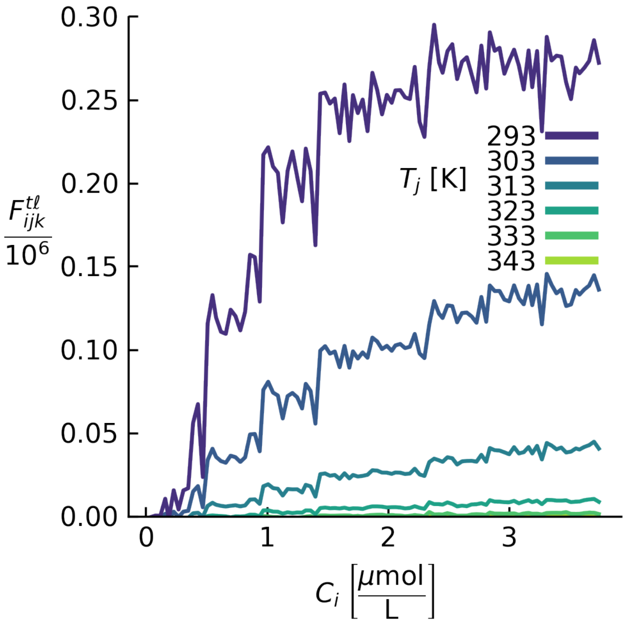


Figure S1: Fluorescence profile associated with dye concentrations given in Table S2.

#### S1.2 Total Least Squares

Equation (22) of the main text may be rewritten as

$$(\mathbf{M}^{\text{TLS}}, \widehat{\mathbf{C}}) = \underset{\substack{\mathbf{M}\in\mathbb{R}^m \\ \mathbf{c}\in\mathbb{R}^n}}{\arg\min} \left\| \mathbf{A} - \mathbf{b}\mathbf{c}^\top \right\|_F^2, \tag{S1}$$

where $\mathbf{A}$ and $\mathbf{b}$ are defined as

$$\mathbf{A} = \begin{bmatrix} \mathbf{F} \\ \rho\mathbf{C}^\top \end{bmatrix}, \qquad \mathbf{b} = \begin{bmatrix} \mathbf{M} \\ \rho \end{bmatrix}.$$

The solution of Equation (S1) is the best rank-1 approximation to $\mathbf{A}$.

To compute the solution, we use an alternating projections algorithm. This algorithm takes advantage of the bilinearity in the objective function, as minimization over one of the variables with the other fixed is a linear least squares problem. Given a guess of $\mathbf{M}^{\mathrm{TLS}}$, denoted as $\mathbf{M}_{-}$, we calculate an updated guess $\mathbf{M}_{+}$ via

$$\mathbf{c}_{+} = \underset{\mathbf{c}\in\mathbb{R}^{n}}{\arg\min} \left\|\mathbf{A} - \mathbf{b}_{-}\mathbf{c}^{\top}\right\|_{F}^{2}, \tag{S2a}$$

$$\mathbf{M}_{+} = \underset{\mathbf{M}\in\mathbb{R}^{m}}{\arg\min} \|\mathbf{F} - \mathbf{M}\mathbf{c}_{+}^{\top}\|_{F}^{2}, \tag{S2b}$$

where $\mathbf{b}_{-} = \begin{bmatrix} \mathbf{M}_{-} \\ \rho \end{bmatrix}$. A natural initial guess is to let $\mathbf{M}_{-} = \mathbf{M}^{\mathrm{LS}}$. To see how much the solution has changed from $\mathbf{M}_{-}$ to $\mathbf{M}_{+}$, we compute the error $\varepsilon = \|\mathbf{M}_{+} - \mathbf{M}_{-}\|$. If $\varepsilon$ is larger than a tolerance value (in this work, we used $10^{-8}$), we assign $\mathbf{M}_{+} \rightarrow \mathbf{M}_{-}$, and calculate an updated $\mathbf{M}_{+}$ via Equation (S2), repeating until $\varepsilon$ is less than or equal to the tolerance. The solutions of Equation (S2) are

$$\mathbf{c}_{+} = \frac{\mathbf{F}^{\top}\mathbf{M}_{-} + \rho^{2}\mathbf{C}}{\mathbf{M}_{-}^{\top}\mathbf{M}_{-} + \rho^{2}}, \tag{S3a}$$

$$\mathbf{M}_{+} = \frac{\mathbf{F}\mathbf{c}_{+}}{\mathbf{c}_{+}^{\top}\mathbf{c}_{+}}. \tag{S3b}$$

Equation (S3a) demonstrates that, $\lim_{\rho\to\infty}\mathbf{c}_{+} = \mathbf{C}$. Substituted into Equation (S3b), this implies that $\lim_{\rho\to\infty}\mathbf{M}_{+} = \mathbf{M}^{\mathrm{LS}}$ (see Equation (21) of main text). In order to account for error in dye concentration, therefore, $\rho$ cannot be large. In this work, we set $\rho^{2} = 0.1$. This choice of weight results in a relatively small error in signal (large error in concentration), as depicted in Figure S2.

The $n+m$ by $n+m$ variance–covariance matrix, $\mathbb{V}$, is calculated via

$$\mathbb{V} = \frac{1}{m(n-1)} \left\|\mathbf{A} - \mathbf{b}^{\mathrm{TLS}}\widehat{\mathbf{C}}^{\top}\right\|_{F}^{2} H^{-1},$$

where

$$\mathbf{b}^{\mathrm{TLS}} = \begin{bmatrix} \mathbf{M}^{\mathrm{TLS}} \\ \rho \end{bmatrix}, \qquad H = \begin{bmatrix} I_{n}(\rho^{2} + \|\mathbf{M}^{\mathrm{TLS}}\|^{2}) & \mathbf{0} \\ \mathbf{0} & I_{m}\|\widehat{\mathbf{C}}\|^{2} \end{bmatrix},$$

and $I_{\ell}$ is an identity matrix of size $\ell$.

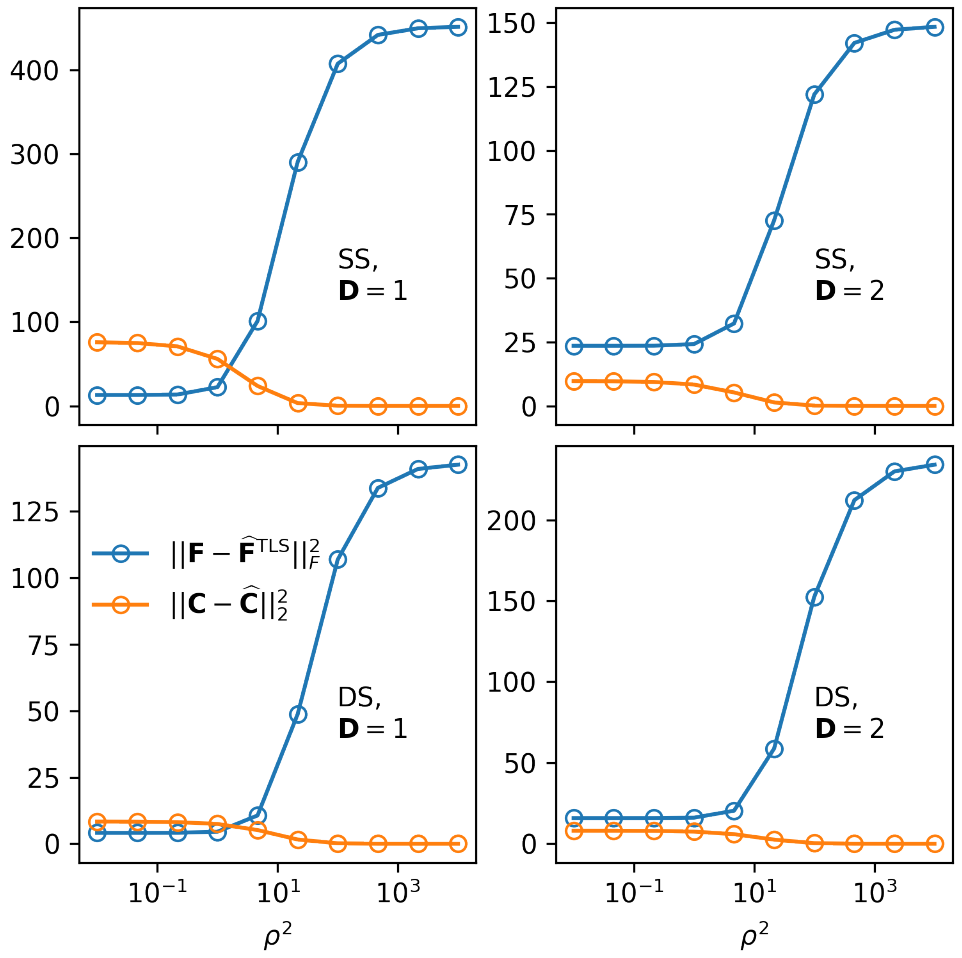


Figure S2: Error in signal and concentration at various values of $\rho$. As $\rho$ increases, the error in fluorescence approaches that of the Least Squares result. When $\rho$ is small, the error in fluorescence is small.

The variance in $\mathbf{M}^{\text{TLS}}$ and $\widehat{\mathbf{C}}$, or $V(\mathbf{M})$ and $V(\widehat{\mathbf{C}})$, respectively, are related to $\mathbb{V}$ as

$$V(\widehat{\mathbf{C}}_i) = \mathbb{V}_{ii}, \qquad i = 1, \dots, n$$

$$V(\mathbf{M}_j) = \mathbb{V}_{jj}, \qquad j = n+1, \dots, n+m$$

### S2 Supplemental Results

#### S2.1 Fraction of Bound Dye

In terms of the scaled and vectorized quantities, defined in Equation (16) of the main text, Equation (6) of the main text becomes

$$\varphi_{j\mathbf{D}} = \frac{\mathbf{K}_j \mathbf{D}}{1 + \mathbf{K}_j \mathbf{D}}, \tag{S5}$$

for a temperature index $j$ and DNA concentration $\mathbf{D}$, where $\varphi_{j\mathbf{D}}$ is defined as $\varphi_{j\mathbf{D}} = c^{\mathrm{b}} \left( \widehat{\mathbf{C}}_i, \mathbf{D}, T_j \right) / \widehat{\mathbf{C}}_i$. Substitution of (25) into Equation (S5) yields

$$\varphi_{j1} = 2r_j - 1, \tag{S6a}$$

$$\varphi_{j2} = 2 - r_j^{-1}, \tag{S6b}$$

where $r_j$ is defined as

$$r_j = \frac{\mathbf{M}_{j1}}{\mathbf{M}_{j2}}. \tag{S7}$$

The dependence of $\varphi_{j\mathbf{D}}$ on temperature $T_j$ and DNA concentration $\mathbf{D}$ is depicted in Figure S3.

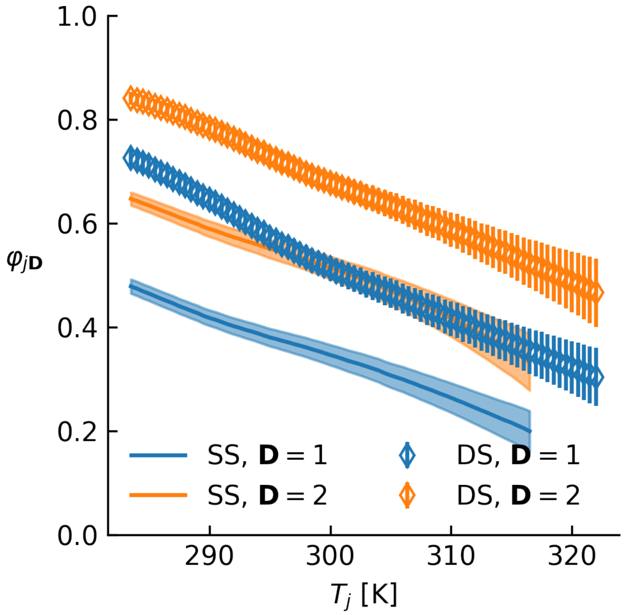


Figure S3: Fraction of dye bound to DNA calculated from Equation (S6) for each temperature $T_j$ and DNA concentration $\mathbf{D}$. The DNA concentration associated with each data set is depicted in its legend entry.

#### S2.2 Average Dye per DNA

In terms of the scaled and vectorized quantities, defined in Equation (16) of the main text, Equation (9) becomes

$$\psi_{ij\mathbf{D}} = \frac{\widehat{\mathbf{C}}_i \mathbf{K}_j}{1 + \mathbf{K}_j \mathbf{D}}, \tag{S8}$$

for a temperature index $j$, DNA concentration $\mathbf{D}$, and well index $i$, where $\psi_{ij\mathbf{D}} = c^{\mathrm{b}}\left(\widehat{\mathbf{C}}_i, \mathbf{D}, T_j\right)/\mathbf{D}$. Substitution of (25) for $\mathbf{K}_j$ into Equation (S8) yields

$$\psi_{ij1} = \widehat{\mathbf{C}}_i \left(2r_j - 1\right) \tag{S9a}$$

$$\psi_{ij2} = \widehat{\mathbf{C}}_i \left(1 - \frac{1}{2r_j}\right) \tag{S9b}$$

where $r_j$ is defined in Equation (S7). The dependence of $\psi_{ij\mathbf{D}}$ on dye concentration $\widehat{\mathbf{C}}_i$, temperature $T_j$, and DNA concentration $\mathbf{D}$ is depicted in Figure S4.

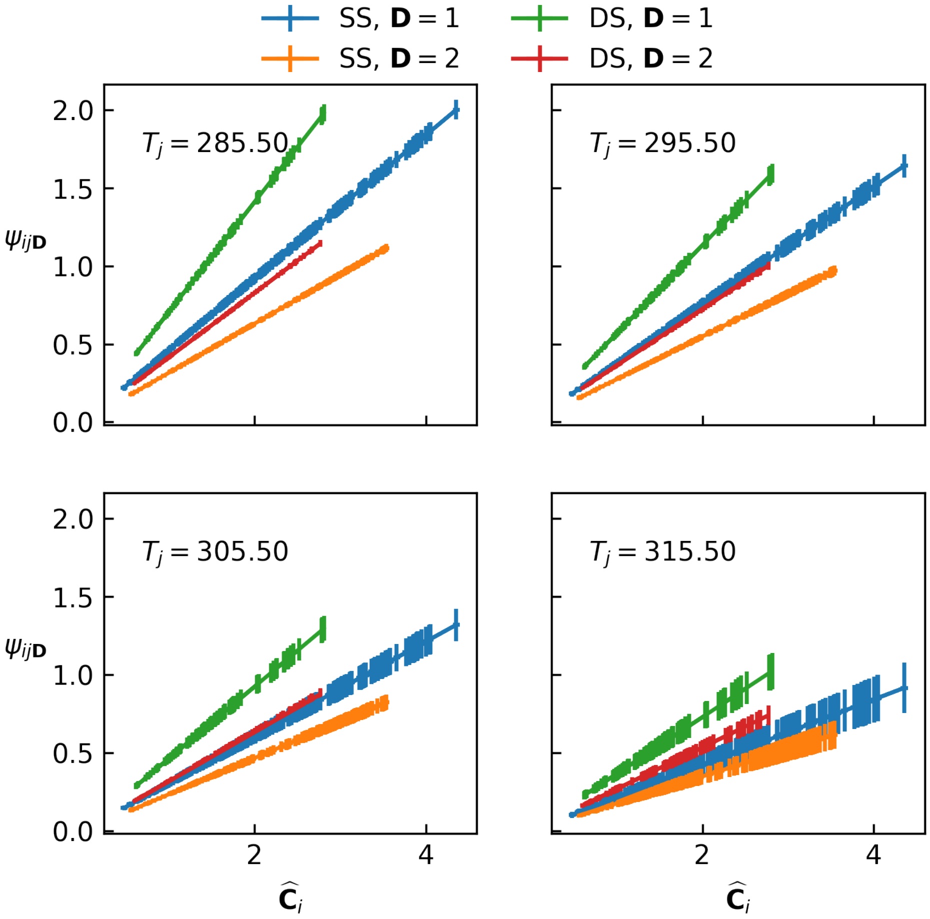


Figure S4: Average number of dye bound per strand of DNA. Calculated from Equation (S9) for each total dye concentration $\widehat{\mathbf{C}}_i$, temperature $T_j$, and DNA concentration $\mathbf{D}$. Different DNA concentrations are depicted in different colored lines. Different temperatures are shown in different subplots.

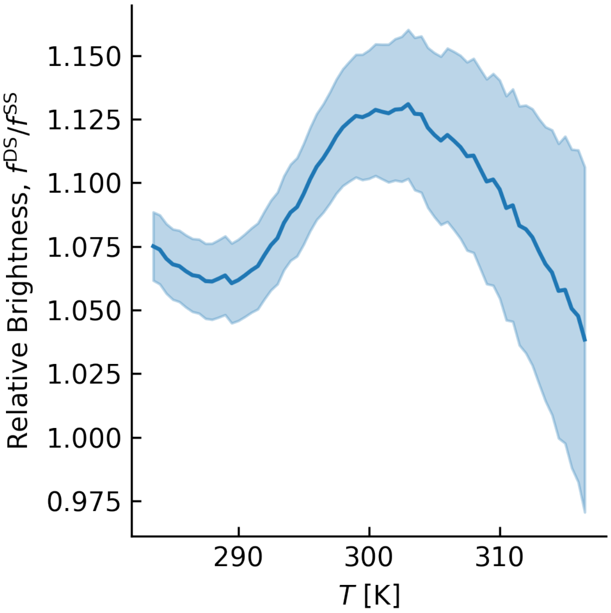


Figure S5: Brightness of double-stranded DNA relative to single-straded DNA, as defined in Equation (28) of the main text, as a function of temperature. Shaded region depicts one standard deviation.
